# Nicotinic modulation of hierarchal inhibitory control over prefrontal cortex resting state dynamics: modeling of genetic modification and schizophreniarelated pathology

**DOI:** 10.1101/301051

**Authors:** Marie Rooy, Fani Koukouli, Uwe Maskos, Boris Gutkin

## Abstract

Nicotinic acetylcholine receptors (nAChRs) strongly modulate the cholinergic drive to a hierarchy of inhibitory neurons in the superficial layers of the PFC, critical to cognitive processes. Genetic deletion of various types of nAChRs, located on specific interneurons, impacts the properties of ultra-slow transitions between high and low activity states (H-states and L-states, respectively), recorded in mice during quiet wakefulness. In addition, recent data indicate that a genetic mutation of the α5 nAChR subunit located on vasoactive intestinal polypeptide (VIP) inhibitory neurons, the rs16969968 single nucleotide polymorphism (α5 SNP), appears to be responsible for “hypofrontality” observed in schizophrenia. Chronic nicotine application to α5 SNP mice restores neural activity to control levels. Using firing rate models of hierarchically organized neural populations, we showed that the change of activity patterns recorded in the genetically modified mice can be explained by a change of activity state stability, differentially modulated by cholinergic inputs to parvalbumin (PV), somatostatin (SOM) or VIP inhibitory populations. A change in amplitude, but not duration of H-states fully account for the lowered pyramidal (PYR) firing rates recorded in α5 SNP mice. We demonstrate that desensitization and upregulation of β2 nAChRs located on SOM interneurons, but not activation of α5 nAChRs located on VIP interneurons, by chronic nicotine application could account for activity normalization recorded in α5 SNP mice. The model implies that subsequent nicotine withdrawal should lead to PYR activity depression more severe than the original hypofrontality caused by SNP mutation.

## Introduction

Alteration of resting activity of the prefrontal cortex (PFC) occurs at the very onset of schizophrenia [1]. Cortical acetylcholine (ACh) release exerts strong modulation of PFC via nicotinic acetylcholine receptors (nAChRs), specifically located on a hierarchically organized circuit of inhibitory neurons within layer II/III [2], making individuals with nAChR gene variants more susceptible to develop mental disorders with cognitive deficits [3]. The α5 SNP, a mutation of the α5 nAChR subunit, is linked to nicotine addiction and also a functional cortical deficit, hypofrontality, a characteristic of schizophrenia patients [4, 5]. Resting-state PFC neural activity in mice with this specific mutation exhibit a reduction, qualitatively similar to the hypofrontality seen in humans and is reversed after 7 days of chronic nicotine application [4].

Neural activity in layers II/III of the mouse PFC during quiet wakefulness is characterized by synchronous ultra-slow fluctuations, with alternating periods of high and low activity [6]. Genetic knock-out (KO) mice of specific nAChRs subunits disrupted these ultra-slow fluctuations in mutant mice, leading to changes in duration of high and low activity states (H-states and L-states, respectively). It has been suggested that slow transitions between activity states could optimize information transmission in the context of lowered metabolism, such as quiet wakefulness, and that it could play an important role in memory consolidation processes [7]. Furthermore, multi-stable dynamics in recurrent networks have been suggested to play a crucial role in working memory and decision-making processes [8, 9]. Interestingly, patterns of hypofrontality in schizophrenia are associated with working memory deficits [10], hypothesized to be a core feature of the disease [11].

Using a circuit computational model, we studied the role of cholinergic inputs to the layer II/III GABAergic interneurons in the modification of PFC synchronous ultra-slow fluctuations, and their impact on the change of neural firing rates in various mice with altered nAChR gene function. We analyzed the effect of chronic nicotine application on specific nAChRs subunits, in order to pinpoint the principal target of nicotine allowing restoration of neural activity in α5 SNP mice to wild-type (WT) levels, supporting the self-medication hypothesis. Our model study of activity pattern alterations in α5 SNP mice lead us to extrapolate that schizophrenia patients may have selective difficulties to gate information rather than to simply retain it. Furthermore, our model was used to predict the consequence of nicotine withdrawal after 7 days of chronic nicotine application on α5SNP mice and highlights the potential dangerous consequences of self-medication in smoking schizophrenia patients.

## Results

### Bistable firing rate dynamics of interconnected neural populations replicates ultraslow fluctuations recorded in the PFC of WT mice

We previously studied experimentally the spontaneous activity of neurons expressing the calcium indicator GCaMP6f in the prelimbic cortex (PrLC) in awake mice by two-photon calcium imaging [4]. You can see in Fig. 1A2-3 an example of a population of simultaneously recorded cells, in a WT mouse, that transit between high activity states (H-states) and low activity states (L-states) lasting several to tens of seconds. We then set out to model the local circuitry that may produce this activity pattern. The circuit model schemed in Fig. 1B1 simulates the firing rate evolution of populations of pyramidal (PYR) neurons connected to a hierarchy of interneurons. Parvalbumin (PV) interneurons, expressing α7 nAChRs subunits [2], target PYR cells axosomaticaly, with strong reciprocal connections [12]. Somatostatin (SOM) interneurons, expressing both α7 and α4β2 nAChRs subunits [2], target the dendrites of PYR cells. The α5α4β2 nAChRs subunits are expressed only by vasoactive intestinal polypeptide (VIP) interneurons [13], that preferentially inhibit the SOM cells, and to a lesser extent PV cells [14]. Both SOM and VIP interneurons receive excitatory feedback from PYR neurons [15, 16]. The model is able to reproduce the ultraslow fluctuations of PYR population activity recorded in WT mice (Fig. 1B2-3) by assuming that two stable states of activity arise from the connectivity between the various population types (see Methods for more information on the model and fitting procedure). Simultaneous network transitions between activity states, for all neural types, are driven by activity fluctuations. This prerequisite is consistent with our experimental findings, showing that the various neuron types have similar H-state and L-state durations (Fig. 1A4, Fig. 1B4 and Supplementary Fig. 8).

**Figure 1:**
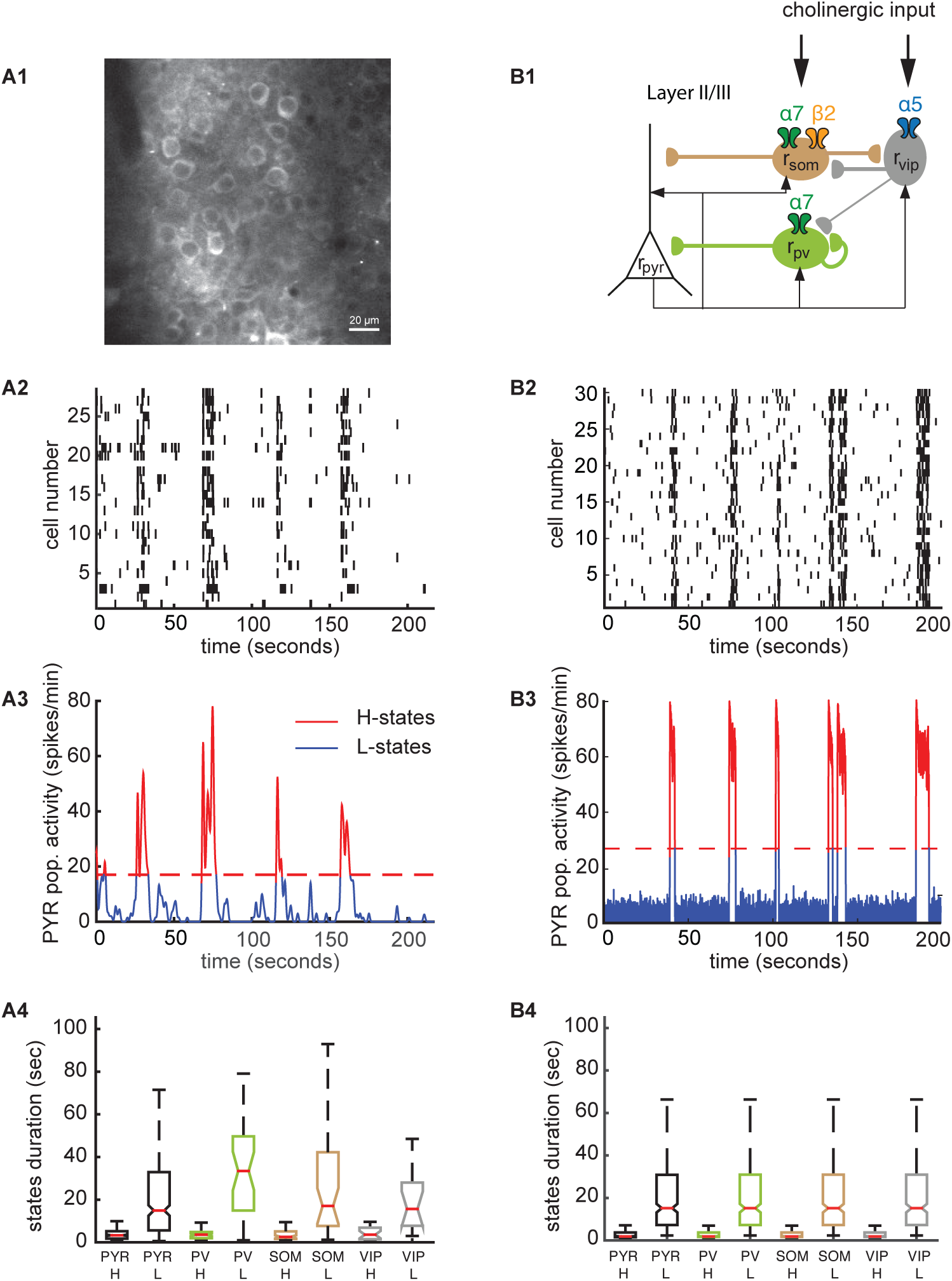
Bistable firing rate dynamics of interconnected neural populations replicates ultraslow fluctuations recorded in the PFC of WT mice. **(A1)** Two-photon image of GCaMP6f expressing neurons, from [4]. (Scale bar: 20 μm). **(A2)** Spike trains of a population of simultaneously recorded cells in a WT mouse, obtained through deconvolution of spontaneous Ca^2+^ transients. 80% of the recorded cells are PYR neurons [4]. **(A3)** Time varying population mean activity of the neurons shown in A2. The dashed red line delineates the threshold between high and low activity states (H-states and L-states, respectively). Red periods correspond to H-states and blue periods to L-states. See [6] for more info on the methods. **(A4)** H-states and L-states durations recorded in the different neuron types. H-states durations are similar between neuron types (mean of ∼3 seconds, no statistical differences), as well as L-states durations (mean of ∼20 seconds, no statistical differences). **(B1)** Schematic of the studied circuitry. **(B2)** Reproduction of single pyramidal neurons activity from the population rate model, using Poisson process with λ(t)=mean population rate(t). **(B3)** Time varying mean population activity of pyramidal neurons, computed from the network model. We use the same method as [6] to delineate H-states (in red) and L-states (in blue). **(B4)** The model reproduces the similar H-states and L-states durations across neural types.

### Effects of changing subtractive vs. divisive inhibitory inputs to the pyramidal population on activity state stability

SOM and PV interneurons activity affect differentially PYR activity through two distinct inhibitory pathways: while SOM preferentially target PYR neurons dendrites, exerting subtractive type of inhibition through an increase of the spiking threshold, PV mainly target the perisomatic region of PYR cells, exerting divisive type of inhibition [17, 18, 19]. Here, we study the effects of subtractive vs. divisive inhibition on bistable network dynamics, without considering PYR neurons excitatory feedback to inhibitory populations, and VIP input to PV population (weaker than the VIP to SOM connection strength [15]). This preliminary analysis permits us to isolate the effects of each type of inhibitory mechanism on PYR activity dynamics, allowing a better understanding of the various phenomena predicted by the fully connected network.

Varying the level of divisive vs. subtractive inhibition of PYR activity leads to changes of activity state stability. Reducing divisive inhibition of PYR activity, for example through a decrease of cholinergic currents to PV interneurons, mimicking the absence of nAChRs (KO), leads to a change of shape for the neuronal input—output function which relates the firing rate to the synaptic input (Fig. 2A1). This modification increases the slope and the maximal firing rate. This in turn leads to a change of stability for both the L-state and the H-state, as evident from the energy landscape of the pyramidal activity (Fig. 2A2): the energy wells of both states become deeper in the KO case. If we assume that transitions between activity states are driven by similar levels of activity noise in the WT and KO conditions, it is easy to see that increasing the stability of both activity states by a decreased level of divisive inhibition makes those transitions less likely. In Fig. 2A3, we can see the bifurcation diagram for the network stable states as a function of PV interneurons external input current. In the range of input values for which the network exhibit bistable dynamics, we notice an increase of durations for the H-states and L-states, consistent with the decrease of transition rate between both states (Fig. 2A4).

**Figure 2:**
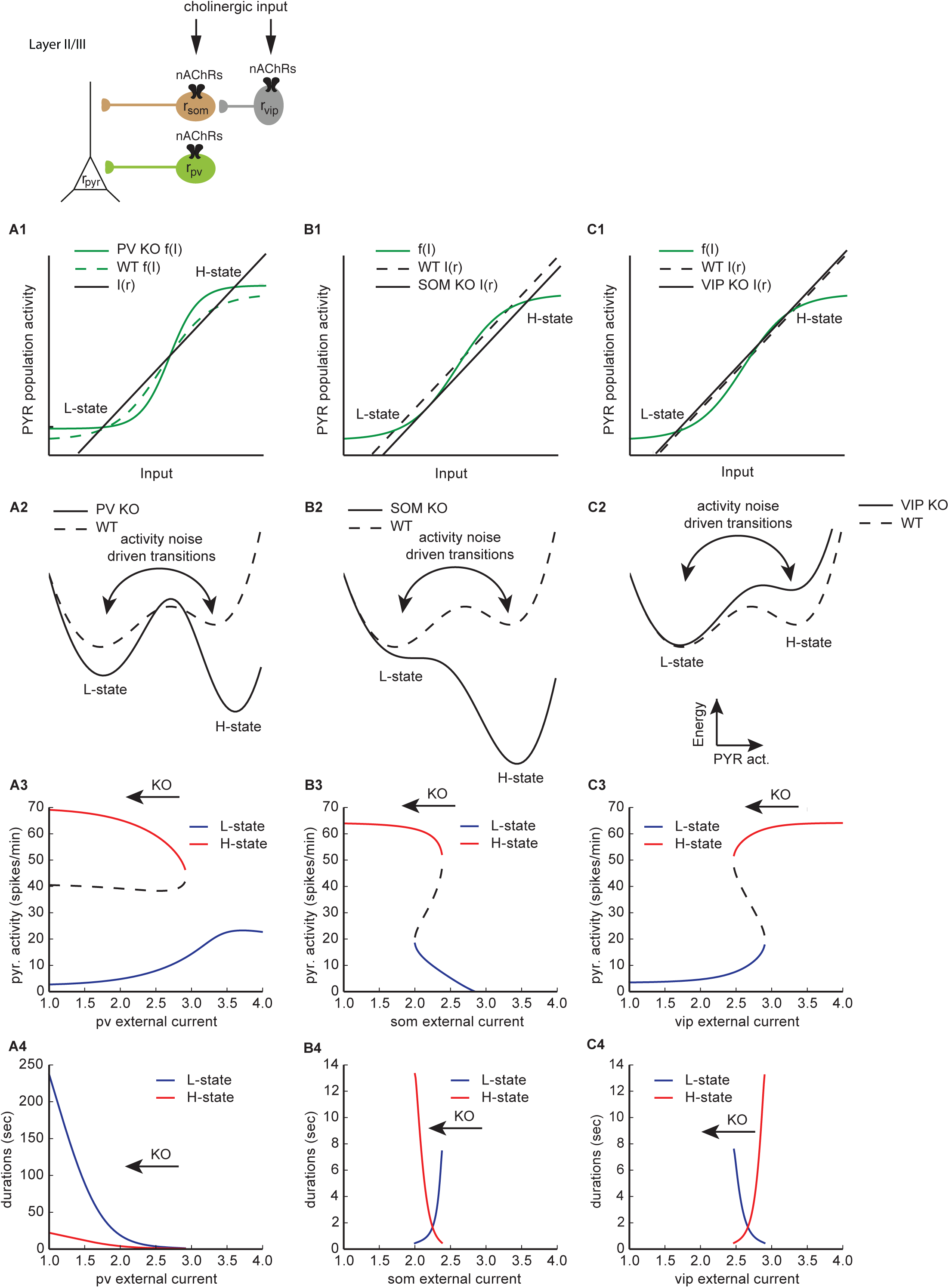
Effects of changing subtractive vs. divisive inhibitory inputs on pyramidal population state stability. PYR excitatory feedbacks to GABAergic neural populations are not considered in this section, nor VIP to PV inputs. **(A1)** Output PYR activity, r, feeds back through recurrent connections and influence synaptic input I(r), which leads to new output firing rates f(I). When r = f(I(r)), two stable states emerge, one corresponding to a high level of PYR activity (H-state), and the other to a low level of PYR activity (L-state), separated by an unstable state. In black, I(r) holds for both WT and PV KO (knock out of nAChRs located on PV population) simulations. Due to divisive inhibition, f(I) changes shape for PV KO simulations (green plain line), compared to WT simulations (green dotted line). **(A2)** Potential energy landscape representation of PYR activity dynamics. Activity fluctuations drive the transitions between H-state and L-state (the two wells), across the unstable state (the hill). The depth of the wells reflects the stability of the activity state. KO of nAChRs on PV (black dotted line) population increases for H-state and L-state stabilities compared to WT (black plain line). **(A3)** H-state (red line) and L-state (blue line) PYR activities as a function of the external input current to PV population. The dashed black line shows the evolution of the unstable activity state. KO of nAChRs is associated with a decrease of external currents (black arrow). **(A4)** Same as (A3), for H-state (red line) and L-state (blue line) durations. **(B1)** For both WT and SOM KO (knock out of nAChRs located on SOM population) simulations, f(I) keeps the same shape (green plain line), whereas I(r) shifts for SOM KO (black plain line), compared to WT (black dotted line), due to subtractive inhibition. **(B2)** Same as (A2), for KO of nAChRs on SOM (black dotted line) population. H-state increase in stability while L-state decrease in stability compared to WT (black plain line). **(B3)** Same as (A3), for SOM external current. **(B4)** Same as (B3), for H-state (red line) and L-state (blue line) durations. **(C1)** For both WT and VIP KO (knock out of nAChRs located on VIP population) simulations, f(I) keeps the same shape (green plain line), whereas I(r) shifts for VIP KO (black plain line), compared to WT (black dotted line), due to subtractive disinhibition. **(C2)** Same as (A2), for KO of nAChRs on VIP (black dotted line) population. H-state decrease in stability while L-state increase in stability compared to WT (black plain line). **(C3)** Same as (A3), for VIP external current. **(C4)** Same as (C3), for H-state (red line) and L-state (blue line) durations.

Reducing subtractive type of inhibition of PYR activity, through a decrease of SOM activity, leads to a shift of the linear relationship between the input current to PYR populations and its firing rate (see Fig. 2B1), such that an asymmetry is created in the change of state stability: the L-state energy well decrease in depth while the H-state energy well increase in depth, in Fig. 2B2. According to the bifurcation diagram in Fig. 2B3, PYR activity should lose its bi-stability when subject to a decrease of external input to SOM population after a critical value. Before this critical point, the H-state should increase in duration while the L-state should decrease in duration (Fig. 2B4). When decreasing the external input to VIP, the linear relationship between the total input current and PYR neurons firing rate shifts in the opposite direction compared to SOM (Fig. 2C1) leading to opposite changes of stability for H-states and L-states (see Fig. 2C2). The bifurcation diagram for stable states along VIP external input current also shows a critical value for bistability (Fig. 2C3), but the change of state durations (Fig. 2C4) is reversed compared to the bifurcation diagram for SOM external input current.

### Effects of changing external inputs to inhibitory populations on network state stability in the fully connected network

When considering the fully connected network, excitatory feedback from pyramidal neurons to the various interneurons subtypes are taken into account, and inhibitory inputs from VIP to PV neurons. Fig. 3A1-3 and Fig. 3B1-3 show that most of the general analysis made for the simplified network holds for the full network, with an expected increase of both H-state and L-state durations for decreased external inputs to PV population (Fig. 3B1), a predicted increase of H-state durations while a decrease of L-state durations for decreased external inputs to SOM population (Fig. 3B2), and a predicted decrease of H-state durations for decreased external inputs to VIP population (Fig. 3B3). The most noticeable difference lies, first, in the change of shape for the bifurcation diagram as function of the external input to PV population (Fig. 3A1): due to high levels of excitatory feedback for PYR to PV population [11], for external inputs higher than a critical value, the network loses its bistability to a single H-state ( as opposed to the single L-state in the simple version of the model, see Fig. 2A3). Second, we observe a slight decrease of L-state durations for decreased external inputs to VIP population (Fig. 3B3), contrary to what was predicted from the simple version of the model (Fig. 2C3). This phenomenon is explained by the modeling of VIP to PV connection: decreased VIP activity leads to increased PV activity, which decreases L-state durations, due to increased divisive inhibition of PYR activity.

**Figure 3:**
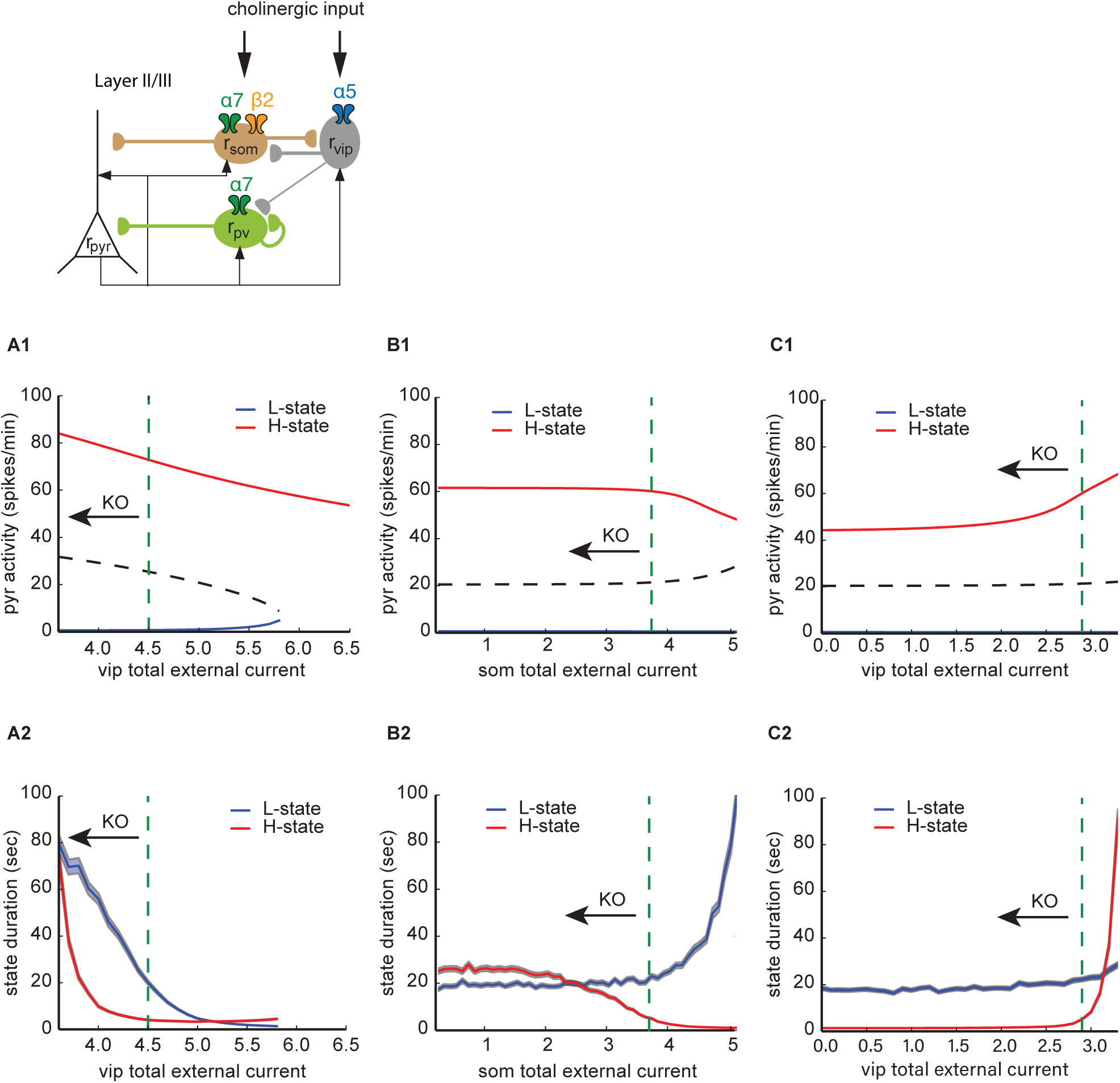
Effects of changing external inputs to inhibitory populations on network state stability in the fully connected network. **(A1)** H-state (red line) and L-state (blue line) PYR activities as a function of the external input current to VIP interneurons population. The dashed green line shows the selected parameter value to reproduce WT mice neural dynamics. KO of nAChRs is associated with a decrease of external currents (black arrow). **(A2)** H-state (red line) and L-state (blue line) durations as a function of the external input current to PV interneurons population. The shaded areas delineate ±sem. The dashed green line shows the selected parameter value to reproduce WT mice neural dynamics. **(B1)** Same as (A1), but for the external input current to SOM interneurons population. **(B2)** Same as (A2), but for the external input current to SOM interneurons population. **(C1)** Same as (A1), but for the external input current to VIP interneurons population. **(C2)** Same as (A2), but for the external input current to VIP interneurons population.

### Accounting for the nAChR KO and mutation impact on ultraslow fluctuations

We explicitly modeled nAChRs levels of activation through [20], in order to have an estimation of the relative effects of β2- and α7-containing nAChRs on the changes of external currents to their target inhibitory neurons. We chose not to take into account desensitization of nAChRs by physiological levels of ACh because of its rapid breakdown through acetylcholinesterase [21, 22]. According to this model, β2 nAChRs, which have high affinity to acetylcholine (ACh), activate such that the underlying cholinergic input is ∼35 folds the amplitude of the cholinergic current due to α7 nAChRs. You can see in Fig. 4A1 (part of the data is from [6]) that experimentally, knocking out α7 nAChRs, located on both PV and SOM interneurons, induce an increase of H-state mean durations compared to WT animals, from 4.2 ± 0.3 to 6.1 ± 0.5 sec, with P<0.05. We could reproduce this effect through modeling, with an increase of mean H-states durations from 4.0 ± 0.2 sec, for simulated WT animals, to 5.6 ± 0.3 sec, for simulated α7 KO animals (P<0.001, see Fig. 4B1), consistent with the effects of divisive and subtractive inhibition on H-state durations, studied in the previous section. Knocking out β2 nAChRs, located on SOM interneurons, induce further increase of H-state durations (8.7 ± 0.5 sec) compared to α7 KO and WT animals. The model accounts for this feature through the modeling of β2 nAChRs high affinity to ACh, with an increase of H-state mean durations to 9.7 ± 0.8 sec. The model predicts lower H-state mean durations for α5 KO (2.0 ± 0.1, P<0.001) compared to WT mice, similar to experimental findings (2.6 ± 0.2 sec, P<0.001), and consistent with α5-containing nAChRs modulatory effects on VIP activity, which inhibit both PV and SOM interneurons. We found no significant changes of H-state mean durations for α5 SNP compared to WT mice, both experimentally and through modeling (4.0 ± 0.3 sec of H-state durations for experiments, and 3.3 ± 0.2 sec for simulations). A significant decrease of mean L-state durations is expected for β2 KO animals, from 22.6 ± 1.3 sec to 13.9 ± 0.8 sec, P<0.01 (see Fig. 4B2), through combined effects of decreased cholinergic input to SOM and VIP populations, as discussed in the previous section, which reproduces experimental findings (21.7 ± 1.3 sec to 15.3 ± 1.0 sec, P<0.001, Fig. 4A2, modified from [6]). No significant change of L-state durations was found for α7, α5 and α5SNP compared to WT, both experimentally and through modeling. We found that 65.9 ± 5.7 *%* of populations exhibited high and low activity states transitions dynamics in β2 KO mice, a significant decrease compared to WT animals (90.3 ± 5.0 % of populations, P<0.05, Fig. 4A3, modified from [6]). According to the model, decrease of cholinergic input to SOM population in β2 KO mice induce an increase of H-state stability while a decrease of L-state stability, such that in some β2 KO mice neural recordings, neurons will spend all of the recorded time in the H-state. The simulations reproduce the decrease of proportion of populations exhibiting L-state/H-state transitions from 88.6 ± 3.6 % in simulated WT animals to 64.3 ± 4.9 % for simulated β2 KO mice (Fig. 4B3). Furthermore, the model predicts that knocking out each type of nAChRs should induce a decrease of H-state amplitudes, and no changes of L-state amplitudes, which is what we found experimentally (see Fig. 4C).

**Figure 4:**
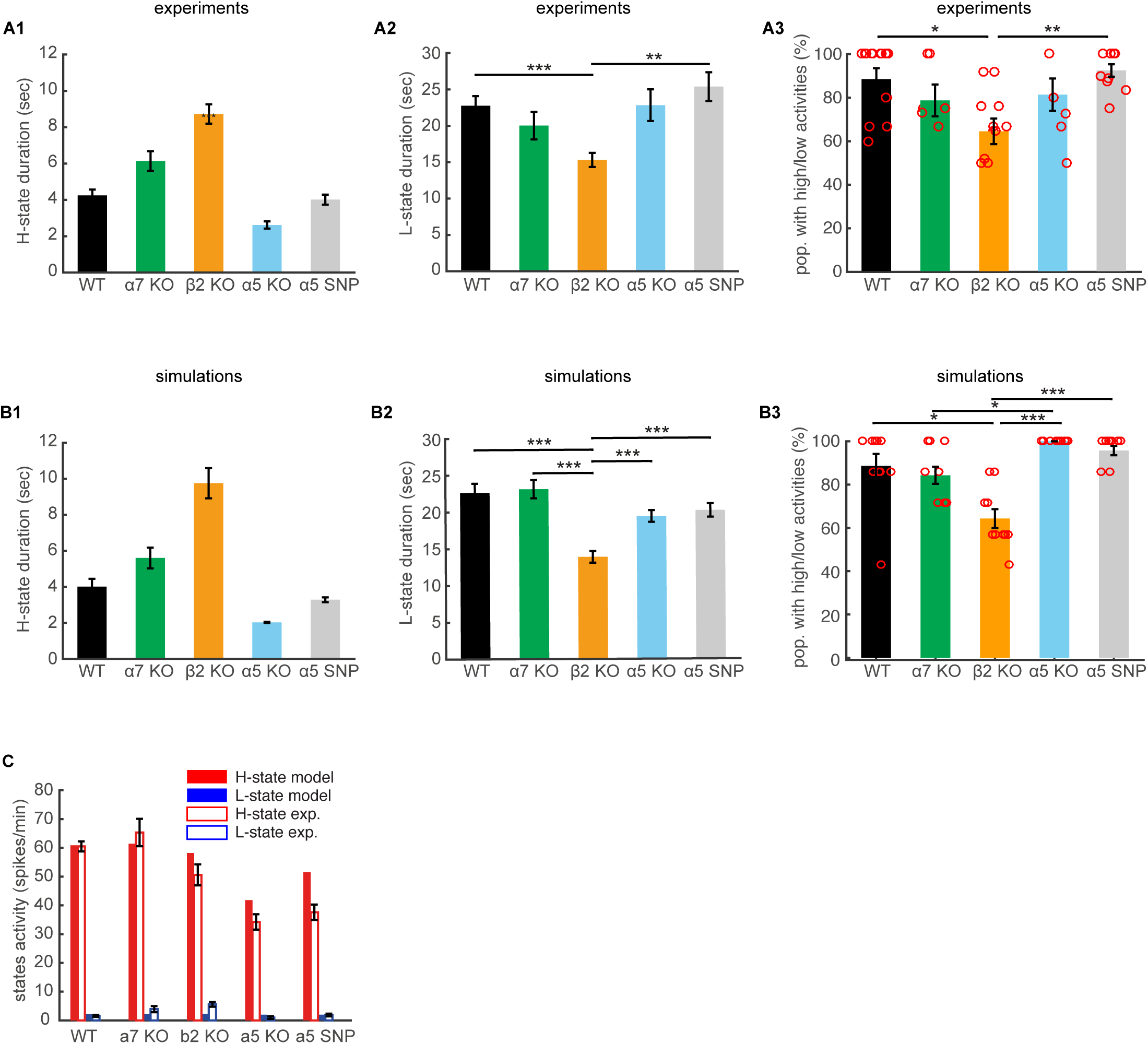
Accounting for the nAChR KO and mutation impact on ultraslow fluctuations. **(A1)** Mean of H-state durations, for WT and mutant mice, modified from Fig. 3C in [6] for α7 KO and β2 KO mice. For α5 KO and α5 SNP mice, we use the same method as in [6]. All mutant mice distributions are significantly different from WT (Kruskal Wallis, P<0.001), except for α5SNP mice. The error bars are ±sem. **(A2)** Mean of L-state durations, for WT and mutant mice, modified from Fig. 3B in [6] for α7 KO and β2 KO mice. β2 KO mice L-state durations are significantly lower compared to WT (Kruskal Wallis, P<0.001). The error bars are ±sem. **(A3)** Mean % of populations with H-states and L-states transitions. The error bars are ±sem. The circle shows the proportions computed for single mice. β2 KO mice exhibit significantly lower % of populations with H-states and L-states transitions (ANOVA, P<0.05), compared to WT mice. Modified from Fig. 3D in [6] for α7 KO and β2 KO mice. **(B1-2-3)** Same as (A1-2-3), computed from simulations. **(C)** H-states (red bars) and L-states (blue bars) activity levels from simulations (filled bars) and experiments (empty bars). Error bars are ±sem. Experimental data described in [6].

### Reproduction of the effects of KO and mutations of nAChRs on neural firing rates

From the modeled population activities, computed for each neuron type and each mouse type, we simulated the firing rate activity of single neurons using a log-normal process (see Methods), and compared the distribution of each mouse type neural activities to experimental findings. Looking at Fig. 1j in [4], we notice that recorded PYR neuron activities in α7 KO (45.9 ± 1.8 spikes/min) and β2 KO (85 ± 1.1 spikes/min) animals increased compared to WT mice (27.5 ± 0.9 spikes/min, P < 0.001 for all comparisons). According to the analysis of activity patterns in WT and KO conditions, increased PYR firing rate activity recorded in mutant mice is expected to be mainly due to increases in the duration of H-states. The high increase of firing rate activities recorded in β2 KO animals is further explained by the decrease in L-state durations. Changes of α7 KO and β2 KO PYR firing rate activities could not be explained by increases in neither H-states nor L-states amplitudes (see Fig. 4C). The model confirms those hypotheses, with an increase of simulated WT activity from 29.0 ± 1.3 spikes/min to 45.0 ± 1.7 spikes/min for α7 KO mice and to 77.8 ± 1.8 spikes/min for β2 KO mice (P < 0.001 for all comparisons, see Fig. 5A). On the other hand, the significant decrease of activities recorded in α5 KO and α5 SNP PYR neurons (8.2 ± 0.5 spikes/min and 20.3 ± 0.6 spikes/min, respectively, P < 0.001 for all comparisons, see Fig. 1f in [4]) are reproduced by the model by both a decrease in H-state durations, and a decrease in H-state amplitudes, with simulated median frequencies of 10.1 ± 0.4 spikes/min for α5 KO mice and of 19.9 ± 0.8 spikes/min for α5 SNP mice (Fig. 5A).

**Figure 5:**
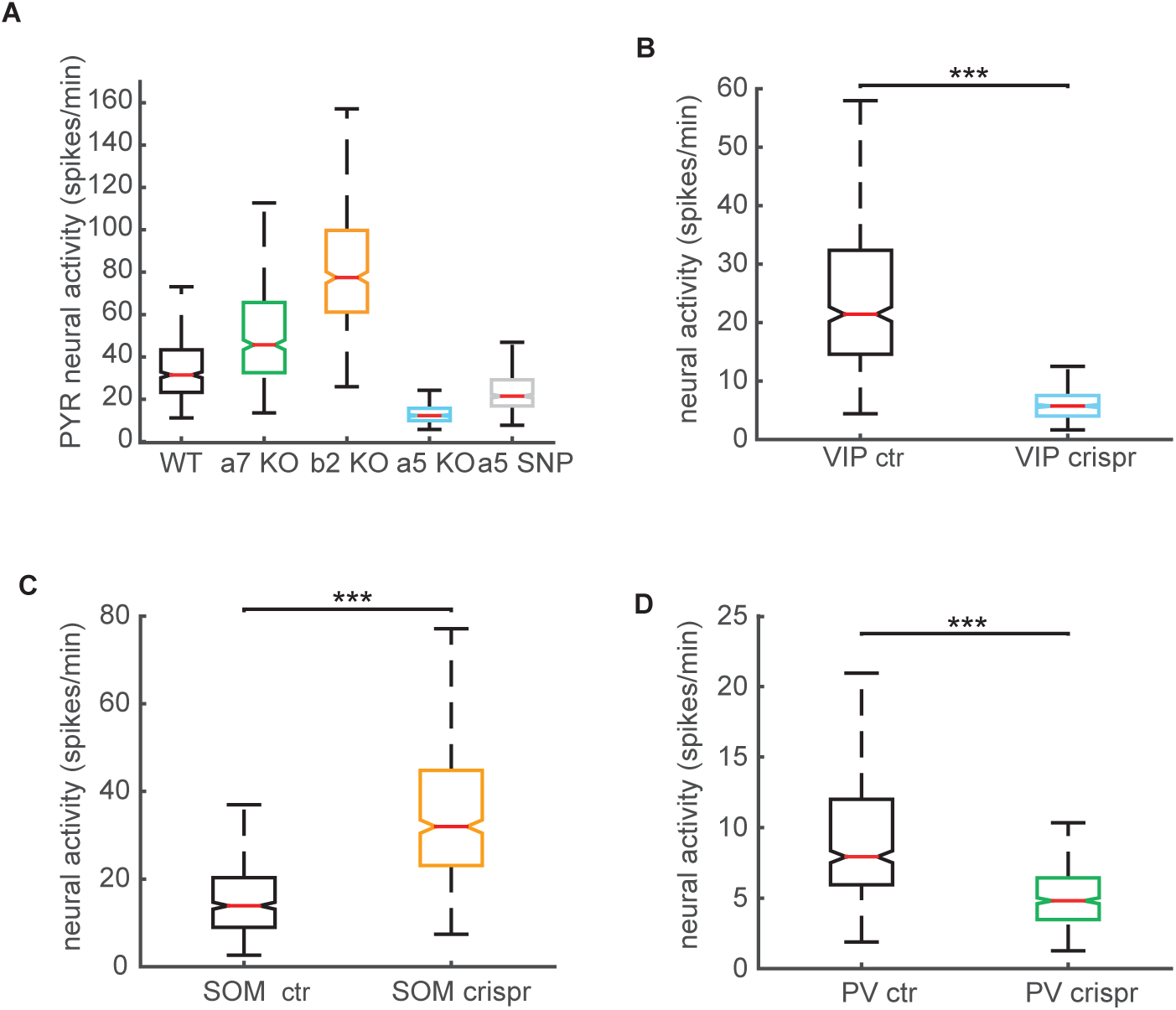
Reproduction of the effects of KO and mutations of nAChRs on neural firing rates. **(A)** Boxplots of PYR neurons firing rates for WT and mutant mice, computed from simlations. All distributions are significantly different (Kruskal Wallis, P<0.001). **(B)** Boxplots of VIP interneurons baseline activities for WT and CRISPR mice. CRIPSPR mice exhibit lower neural activities compared to WT mice (Kruskal Wallis, P<0.001). **(C)** Boxplots of SOM interneurons baseline activities for WT and CRISPR simulated mice. CRIPSPR mice exhibit higher neural activities compared to WT mice (Kruskal Wallis, P<0.001). **(D)** Boxplots of PV interneurons baseline activities for WT and mutant mice affected by the CRISPR technology for the deletion of the α5 subunits, according to simulations. CRIPSPR mice exhibit similar levels of neural activity compared to WT mice.

To further confirm our hypothesis that decreased disinhibition in α5 KO and α5 SNP mice accounts for hypofrontality, we compared the changes of VIP, PV, and SOM interneurons under α5 knock-down in experiments and simulations. Clustered, regularly interspaced, short palindromic repeats (CRISPR)-associated endonuclease (Cas)9 technology were used to knockdown the α5 subunits *in vivo*, as shown in [4]. It has been found experimentally that VIP neurons median activity changed under the CRISPR technology from 21.5 ± 2.17 spikes/min to 3.5 ± 0.7 spikes/min (P<0.001, see Fig. 3f in [4]). We could account for this significant decrease in activity in the model by decreased H-state network durations and decreased VIP H-state activity (see Fig. 4A1, Fig. 4B1 and Supplementary Fig. 9C3). As a result, the simulated spike frequency of VIP interneurons in VIP^CRISPR^ mice (5.7 ± 0.2 spikes/min) was significantly lower than VIP neural activity in simulated WT animals (21.4 ± 1.2 spikes/min, P<0.001, see Fig. 5B). The model predicted that the decreased VIP levels of activity should result in increased levels of SOM activity according to the disinhibition hypothesis. Experimental findings endorsed this prediction, through a robust increase in SOM interneuron spontaneous activity (39.12 ± 3.14 spikes/min) compared to control mice (5.57 ± 1.31 spikes/min, P<0.001, see Fig. 3n in [4]). The model could reproduce this increase of SOM activity despite the significant decrease of network H-state duration (Fig. 4A1 and Fig. 4B1) through an increase in SOM H-state level of activity (Supplementary Fig. 9C2). SOM WT mice simulated median activity increased from 13.9 ± 0.7 spikes/min to 32.0 ± 1.2 spikes/min (P<0.001, see Fig. 5C). Experimentally, the decrease in PV^CRISPR^ interneuron activity was not significant (6.07 ± 0.60 spikes/min) compared to control mice (5.62 ± 0.45 spikes/min, see Fig. 3j in [4]). This feature poses a challenge to the layer II/III model. Because PV interneurons receive high levels of excitatory input from PYR neurons, we would expect a decrease of excitatory input to this population in α5 KO and CRISPR mice, due to decreased PYR neurons firing rate. This decreased excitatory input may be compensated by a decreased inhibition from the less active VIP interneurons, yet this connection is rather weak. We confirmed this in the model (Fig. 5D), we saw a slight but significant decrease of PV^CRISPR^ activity compared to PV activity in WT simulated mice.

### Desensitization and upregulation of β2 nAChRs normalizes α5 SNP mice network activity to WT levels after chronic nicotine application

Based on the evidence that nicotine administration to α5 SNP mice by mini-pump infusion increased PYR neuron activity to WT levels in the PFC of mice [4], indicating that it could reduce some of the cognitive deficits linked to schizophrenia, we used our model to pinpoint the specific nAChRs responsible for this normalization. We know that β2-dependent nAChR currents, but not α7, innervating interneurons in layer II/III of PFC, completely desensitize after exposure to smoking concentrations of nicotine in slice preparation [23], with an exception for α5α4β2 nAChRs, that are more resistant to desensitization [24]. Modeling nAChRs levels of activation and desensitization [20] in contact of physiologically realistic levels of nicotine during smoking permits to predict the exact change of cholinergic currents amplitude, for each specific interneuron subtype. The model predicts activations of α7 and α5-containing nAChRs in contact of 1 μM of nicotine and a strong desensitization of β2-containing nAChRs (see table 6 and Fig. 6A1). As a result, you can see in Fig. 6A1 and Fig. 6A2 our predictions for PYR activity variations in WT and α5 SNP mice when nicotine targets selectively each type of nicotinic receptor. Desensitization of β2 nAChRs decreases cholinergic inputs to SOM interneurons, and should increase the H-state network duration, leading to higher PYR firing rates in both WT and α5 SNP animals. An increase of cholinergic inputs to both SOM and PV interneurons, through the activation of α7 nAChRs, is assumed to induce a decrease of H-state durations, reducing PYR activity in WT and α5 SNP mice. Activation of α5 nAChRs, increasing cholinergic inputs to VIP interneurons, should disinhibit PYR neurons, causing higher PYR activities in both WT and α5 SNP mice. As a consequence, in α5 SNP mice, nicotine application leads to higher PYR activities compared to WT mice exclusively through its interaction with β2 nAChRs (Fig. 6A2). Looking at the experimental results (Fig. 4c in [4]) we see that in both WT mice and α5 SNP mice, nicotine induces high increase of PYR activity after two days of nicotine administration. The simulation of nicotine effects on layer II/III nAChRs permits to replicate those results (see Fig. 6B), and we know from our preliminary analysis in Fig. 6A2 and Fig. 6A3 that those effects are almost entirely due to the desensitization of β2 nAChRs.

**Figure 6:**
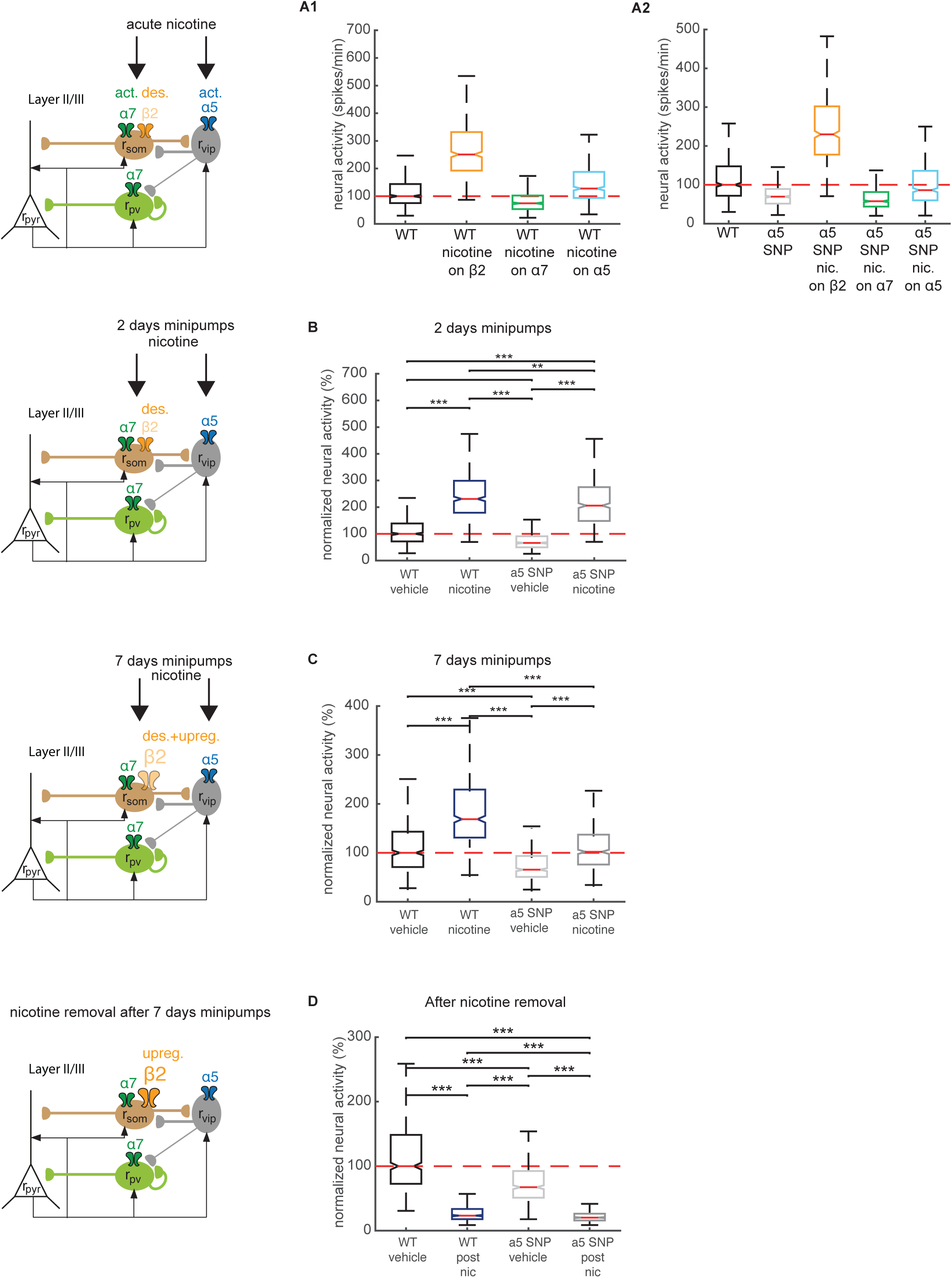
Desensitization and upregulation of β2 nAChRs normalizes α5 SNP mice network activity to WT levels after chronic nicotine application. **(A1)** Distribution of PYR neurons firing rates for WT mice, computed from simulations of nicotine effects on α7, β2, and α5 nAChRs. All distributions are significantly different (Kruskal Wallis, P<0.001). **(A2)** Distribution of PYR neurons firing rates for WT and α5SNP mice, computed from simulations of nicotine effects on α7, β2, and α5 nAChRs in α5SNP mice. All distributions are significantly different (Kruskal Wallis, P<0.001). **(B)** Distribution of PYR neurons firing rates for WT and α5SNP mice, control and treated with 2 days of chronic nicotine application, obtained from simulations. All distributions are significantly different (Kruskal Wallis, P<0.001). **(C)** Distribution of PYR neurons firing rates for WT and α5SNP simulated mice, control and treated with 7 days of chronic nicotine application. All distributions are significantly different (Kruskal Wallis, P<0.001). The model predicts an upregulation of β2 nAChRs. **(D)** Distribution of PYR neurons firing rates for WT and α5SNP mice, control and after nicotine removal following 7 days of chronic nicotine administration, predicted from simulations. All distributions are significantly different (Kruskal Wallis, P<0.001).

We can notice in Fig. 4d, in [4], that the increase in PYR activity after 7 days of nicotine administration is reduced in both WT mice and α5 SNP mice compared to the 2 days treatment, such that PYR neurons firing rate in α5 SNP mice treated with nicotine are at the level of WT mice. Previous studies have indicated that long-term nicotine exposure over days increases or upregulates the number of high-affinity nicotine binding sites on α4β2 nAChRs [25]. In addition to this, no nicotine-induced upregulation was observed for α5α4β2 nAChRs [26]. Hence, we tested through modeling the effect of the upregulation of β2 nAChRs, located on SOM interneurons, on the PYR neuron firing rates. We were able to reproduce the normalization of PYR activity to WT levels in α5 SNP mice after 7 days of administration by considering a 1.8-fold increase of the number of β2 nAChRs located on SOM interneurons (see figure 6C). You can see in Supplementary Fig. 10A1 that the increase of PYR activity in WT animals after 7 days of nicotine treatment is accompanied by an increase of H-state durations, that is reproduced in simulations (Supplementary Fig. 10A2). It is also accompanied by a significant decrease of SOM activity (Fig. 4h in [4]), consistent with the model predictions (Supplementary Fig. 10B).

We used our modeling framework, validated for its consistency with both the KO mice and nicotine application neural recordings, to test the effects of nicotine withdrawal after 7 days of nicotine application. We predict that nicotine withdrawal would rapidly resensitize β2-containing nAChRs, but that the renormalization of the number of receptors, which had been increased through upregulation, would occur on much slower time scales. As a result, SOM interneurons would end up with higher levels of cholinergic innervation in the post-compared to the pre-treatment condition. Both WT and α5 SNP mice would have much lower levels of PYR activity compared to their initial state (Fig. 5D).

## Discussion

### Summary of results

What we learned from this modeling framework is threefold.

First, the change of cholinergic input to the various GABAergic neurons, due to the knockout of various types of nAChRs, could fully account for the change of activity patterns recorded in the various mouse lines. The KO of β2 nAChRs, high affinity receptors to ACh, decreased cholinergic input to SOM population, decreasing subtractive inhibition of PYR activity. Decreasing this specific type of inhibition of PYR activity increases the stability of the network high activity states (H-sates), increasing their durations, plus it decreases the stability of the network low activity states (L-states), decreasing their durations. KO of α5 nAChRs, β2-containing nAChRs, decreases cholinergic inputs to VIP population, decreasing synaptic inhibition of SOM interneurons, thus decreasing subtractive inhibition of PYR activity. It leads to a significant decrease of H-state stability, decreasing their durations, but doesn’t lead to a significant increase of L-states durations, because of VIP to PV decreased inhibitory input, having an opposite effect on L-state stability, through an increase of divisive inhibition of PYR activity. The α7 nAChRs, having low affinity properties to ACh, leads to lower decrease of cholinergic inputs to SOM and PV interneurons, which both, through subtractive and divisive inhibition respectively, increase the stability, thus durations, of H-states.

Second, the predicted changes of H-states and L-states properties caused by KO of nAChRs were sufficient to explain the changes of single neurons firing rates, given that neural activities in a given population had a log-normal, skewed distribution with equal mean and variance (see Methods). Hence, increases of H-state durations, but not activity levels, fully accounted for the increase of PYR neural firing rates in both α7 and β2 KO mice, while both a decrease in H-state durations and levels of activity explained the decreased PYR neural activities in α5 KO and α5SNP mice. As a challenge for the layer II/III model, we were not able to reproduce the invariance of PV interneuron activity after the KO of α5 nAChRs. We posit that considering layer I interneurons, targeted by layer II/III SOM interneurons and targeting layer II/III PV interneurons, may redress this difficulty.

Third, nicotine could restore PYR activity to WT levels in α5SNP mice because of mixed desensitization and upregulation of β2 nAChRs after 7 days of chronic application. Activation of α5 nAChRs by nicotine was not sufficient to compensate α5SNP mice activity deficits. Upregulation of β2 nAChRs after 7 days of treatment should lead to activity depression in both WT and α5SNP mice when nicotine is removed, which could represent a highly critical situation in schizophrenia patients. Hence, this work highlights the potentially deleterious consequences of chronic nicotine taking/withdrawal cycle in schizophrenia, prevalent in this patient population and the risk of increasing the severity of symptoms rather than ameliorating them.

### Cognitive functions of high and low activity states transitions

Transitions between the high and low persistent states of activity have been proposed to play a number of computational roles in processes such as decision making [9] and working memory (WM) [8, 9]. In such attractor-based framework, the firing rate of populations of recurrently connected pyramidal neurons, together with the local inhibitory circuitry, encode information (stimulus characteristics, motor commands etc) that can be maintained actively in working memory through stable states of the activity. The more stable is a memory state, the less susceptible it is to be erased by internal noise or through competition with other potentially irrelevant distractors. Since we know that prefrontal cortex is crucially involved in both cognitive processes [27, 28], studying the change of persistence capabilities of its neural networks under altered cholinergic drive to the various inhibitory populations, allows us to pinpoint the characteristics and the origins of cognitive disruption seen in the corresponding psychiatric disease. Once these are identified, they may be suggested to be targets for an eventual therapeutic intervention. For instance, we know that hypofrontality in schizophrenia is associated with working memory deficits [10], yet one may ask what specific aspects of WM function are affected by the hypofrontality? In our analysis and experiments we did not see any significant change of the high/low activity states stability in α5SNP mice. This result suggests that the stability of the corresponding memory states should not be changed in the α5SNP versus the WT prefrontal-cortex. On the other hand, we found that the H-states in α5SNP mice had lower firing rates. According to previous studies, this may lead to a decreasing information transmission efficacy [7]. Altered information transmission may in turn have an indirect effect on working memory performance, such that relevant stimuli may have a lower efficacy to recruit pyramidal cell populations coding the relevant information to be maintained. Or in other words, the gating of working memories would be disrupted. Hence, we extrapolate that the lower WM capacity in schizophrenic patients with the SNP mutation may be due to encoding or gating impairments, rather than an inefficacy to maintain information in the delay period.

Ultra-slow oscillations seen during quiet wakefulness [30, 31, 32, 33] have also been shown to play a role in memory consolidation by controlling the flow of memory-related information from thalamus and hippocampus to the cortex [34, 35]. Notably, the since transmission of such information appears to be proportional to PYR firing rates [36], the altered H-state activity in α5SNP PFC may imply a memory-consolidation pathology in α5-mutation schizophrenic population.

### Nicotine withdrawal in schizophrenia

Our work further suggests a neurobiological explanation for the high prevalence of smoking and low smoking cessation rate observed among individuals with schizophrenia [37]. Our results show that nicotine cessation should decrease prefrontal activity in both WT and α5SNP PFC, with the most drastic hypofrontality seen in α5SNP mice under nicotine removal. This predicted decrease of pyramidal activity is due to the upregulation of β2 nAChRs located on SOM interneurons, induced by several days of chronic nicotine application. We predict that the lower cessation rates seen in schizophrenics are caused by a pronounced hypofrontality, induced by a combination of mutated α5 nAChRs located on VIP interneurons and the upregulation of β2 nAChRs located on SOM interneurons. In fact, previous work showed that negative affect, one aspect of the negative schizophrenia symptoms associated with hypofrontality, is a key contributor to the low quitting rate seen in smoking schizophrenics [38]. At the same time pharmacotherapy, through the use of varenicline, having similar nAChRs interaction mechanisms to nicotine, increases the abstinence rate in smokers and even more drastically in schizophrenic smokers (from 4.1% to 23.2%) [37]. We may suggest that these studies lend support to our conjecture.

In summary our circuit-based dynamic modelling approach opens a number of further avenues to both study specific disease-related alteration of nicotinic modulation in cortical circuits and to identify potential points-of-entry for therapeutic interventions.

## Methods

Model variables described the dynamics of the firing rates of the various neuronal populations (re, rp, rs, and rv, for PYR, PV, SOM and VIP neurons, respectively) in a local PFC circuit, using a generalization of the Wilson-Cowan model [39], based on [40] theoretical work for subtractive vs. divisive inhibition of PYR activity by SOM and PV interneuron populations respectively. We incorporated subtractive inhibition between SOM, VIP and PV populations [39].

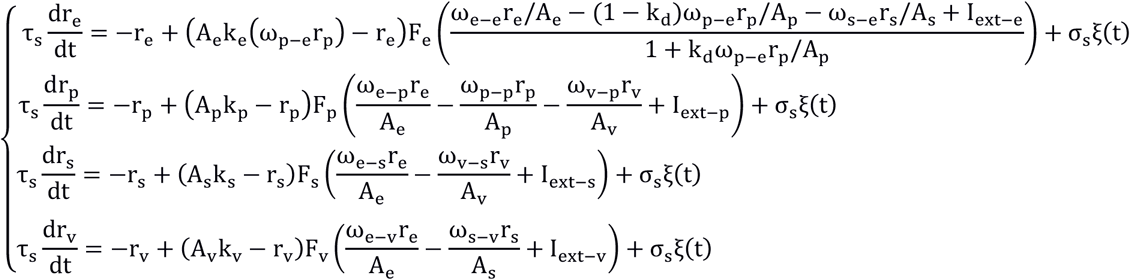

With I_ext-p_ = I_0-p_ + I_α7-p_, I_ext-p_ = I_0-p_ + I_α7-p_ + I_β2-p_, I_ext-v_= I_0-p_ + I_α5-v_. We Set τs = 20 ms, close to each populations’ type membrane time constant [41]. F_X_ is a sigmoid response function characteristic of an excitatory (inhibitory) population, which gives a nonlinear relationship between input currents to a population, and its output firing rate. k_X_ modulate the amplitude of the firing rate response to an input current, dependent on PV activity for PYR neurons. See [39] for the details of F_X_ and k_X_ functions, which are set by two constants, Θ_X_ (minimum displacement) and α_x_ (maximum slope). Most of PV interneurons are Basket cells, thus have a lower input resistance compared to PYR neurons [42]. On the other hand, SOM interneurons, mostly Martinotti cells, and VIP interneurons, have higher input resistance compared to PYR neurons [42]. The highest input resistances are recorded in VIP interneurons [42]. Hence, we changed the maximal slopes α_X_ according to those experimental findings (see table 1). We modelled PV modulation of PYR activity as a combination of both divisive and subtractive inhibition, which can be thought as more biologically realistic [17, 19]. An additional constant parameter k_d_=0.8 is introduced in the model in order to express the fraction of divisive modulation that is delivered to the excitatory population. The rest of the modulation, 1-k_d_, is delivered as subtractive. The ω_X–X_ (with x=e, p) are the self-excitatory (or self-inhibitory) synaptic coupling of the excitatory (inhibitory) neural populations. The ω_X–Y_ (with x≠y, x=e, p, s or v and y= e, p, s or v) are the excitatory (or inhibitory) synaptic coupling from one population to another. We did not consider self-inhibition in the SOM and VIP interneuron populations, since inhibitory chemical synapses between those neurons are rarely observed [41, 43], and did not study PV and SOM direct connections in this version of the model, since regional discrepancies have been reported [41, 44, 45, 46]. The parameter σ_s_ controls the strength of fluctuations of neural populations’ firing rate, and ξ(t) is a Gaussian white noise with mean 0 and variance 1. In the original version of the model [39], activities of the various neural populations were normalized in the range 0 to 0.5. We added factors (A_e_, A_p_, A_s_, A_v_) in order to fit the range of activities to experimental values.

**Table 1:** Input-output functions parameters for PYR, PV, SOM and VIP populations (e, p, s and v).

| $\Theta_e$ | $\alpha_e$ | $\Theta_p$ | $\alpha_p$ | $\Theta_s$ | $\alpha_x$ | $\Theta_v$ | $\alpha_x$ |
| --- | --- | --- | --- | --- | --- | --- | --- |
| 7.0 | 1.9 | 7.0 | 2.6 | 7.0 | 1.5 | 7.0 | 1.2 |

In order to model spike frequency adaptation from PYR neurons in the network model, we used the following equation, adapted from [47]:

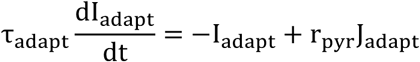

τ_adapt_ and J_adapt_ are the adaptation time constant and the adaptation strength of PYR population. We chose τ_adapt_ = 600 ms according to [48].

## Fitting procedure

We created an optimization algorithm in order to select the parameters (ω_X–X_, ω_X–Y_, J_adapt_ and I_ext–X_) that best reproduced the network dynamics observed in WT animals, for the different neural populations in layer II/III. This algorithm created 10^5^ random associations between parameter values, in the range 1-55 for synaptic connections, and 0.1 to 0.55 for external inputs. VIP to PV connection was set to be half of VIP to SOM connection [14]. For each set of parameters, we computed the roots of the non-linear set of equations 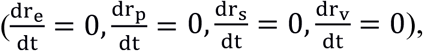 using MINPACK’s hybrd and hybrj algorithms.

First, we selected the parameter sets that displayed two roots, hence two stable states of activity. They represent ∼6% of the total number of tested associations. In supplementary figure 1, you can see the distribution of parameter values for networks that exhibit bistability. First, we see that, for most parameters, there is a uniform distribution of values along the full range of parameters leading to bistability. The noticeable exceptions are the distributions of PYR to PYR connections, which is shifted to higher values, and the distribution of VIP to SOM and SOM to VIP connections, which are shifted to lower values. This is consistent with the reported crucial role of recurrent excitatory connections for sustained high levels of activity [49]. 93%, 61% and 96% of networks exhibited a higher PV, SOM and VIP activity, respectively, in PYR high activity state (H-state) compared to the low activity state (L-state).

Second, we selected connection sets for which the normalized H-state is far from saturation (<0.45), and with a higher activity in H-state than L-state for all neuron types, so that the transitions between H-states and L-states are simultaneous across cell types. Third, we fitted the factors A_e_, A_p_, A_s_, A_v_ so that the H-state for each simulated population type correspond to the ones found experimentally, computed as the median of all populations of simultaneously recorded cells levels of high activities. We selected the networks for which the error of the L-states levels between the model and experiments, computed as for high activities, didn’t exceed 5 spikes/min, for each neural population type. From those selected networks, we simulated the firing rate time evolution of all neural types, in order to compute the distribution of H-state and L-state (500 states each) durations for varying levels of noise σ_s_, between 0.001 and 0.02. For each selected association of parameters, we selected the noise level that best reproduced the mean and mode of H-states and L-states durations. In a following step, we selected networks with a mean relative error, for the properties of ultraslow fluctuations extracted from the experimental data (see table 2) in WT animals that is below 100%. At this step of the selection procedure, we are left with 13 parameter set candidates, out of the initial 10^5^ networks. You can see in supplementary figure 2 A-D the distribution of the synaptic weights to the various neural populations. If you compare those distributions to the ones in supplementary figure 1, corresponding to all bistable networks, you can notice that there is not a large reduction of the range of selected parameters in the parameter space, despite a large reduction of the number of networks, from 5794 to 13, able to reproduce closely the properties of activity patterns observed experimentally in WT animals. In order to discriminate between the 13 last parameter set candidates, we looked at the change of H-state levels between WT and α5 KO. We choose the network in which change of SOM high activity state closely resembled the one observed experimentally (see supplementary figures 2E and 2F). We used hand fitting (+−5%) on the most sensitive parameters in order to further diminish the error associated to this parameter set. See tables 3–4–5 for all selected parameters.

**Table 2:** Experimental properties used for model fitting.

|  |  |  |  |
| --- | --- | --- | --- |
| PYR L-state median activity (spikes/min) | PV L-state median activity (spikes/min) | SOM L-state median activity (spikes/min) | VIP L-state median activity (spikes/min) |
| 1.75 | 1.04 | 1.12 | 1.33 |
| PYR H-state median activity (spikes/min) | PV H-state median activity (spikes/min) | SOM H-state median activity (spikes/min) | VIP H-state median activity (spikes/min) |
| 60.19 | 35.3 | 35.2 | 68.8 |
| H-state mean duration (sec) | H-state mode duration (sec) | L-state mean duration (sec) |  |
| 4.4 | 1.5 | 21.7 |  |

**Table 3:** Selected synaptic weights, and adaptation strength (arbitrary unit), divided by their corresponding scaling factor A_e_, A_p_, A_s_, A_C_ for PYR, PV, SOM and VIP populations (e, p, s and v).

|  |  |  |  |  |  |  |  |
| --- | --- | --- | --- | --- | --- | --- | --- |
| $\omega_{e-e}/A_e$ | $\omega_{p-e}/A_p$ | $\omega_{s-e}/A_s$ | $\omega_{e-p}/A_e$ | $\omega_{p-p}/A_p$ | $\omega_{v-p}/A_v$ | $\omega_{s-p}/A_s$ | $\omega_{e-s}/A_e$ |
| 0.137 | 0.101 | 0.002 | 0.113 | 0.093 | 0.024 | 0.004 | 0.077 |
| $\omega_{v-s}/A_v$ | $\omega_{e-v}/A_e$ | $\omega_{s-v}/A_s$ | $J_{adapt}/A_e$ | $I_{ext-e}$ | $I_{ext-p}$ | $I_{ext-s}$ | $I_{ext-v}$ |
| 0.048 | 0.042 | 0.001 | 0.056 | 3.9 | 4.5 | 3.6 | 2.9 |

**Table 4:** Fitted scaling factors for realistic ranges of activity for PYR, PV, SOM and VIP populations (e, p, s and v).

| $A_e$ | $A_p$ | $A_s$ | $A_v$ |
| --- | --- | --- | --- |
| 172 | 261 | 757 | 664 |

**Table 5:**
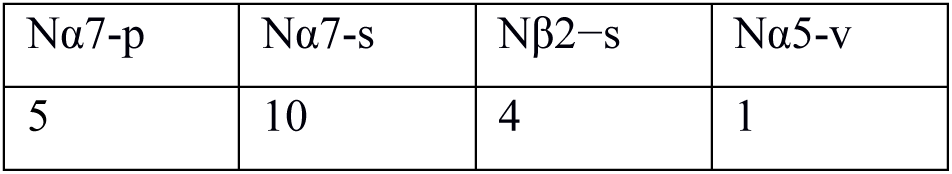
Predicted proportion of nAChRs on each interneuron type.

| $N_{\alpha 7-p}$ | $N_{\alpha 7-s}$ | $N_{\beta 2-s}$ | $N_{\alpha 5-v}$ |
| --- | --- | --- | --- |
| 5 | 10 | 4 | 1 |

In supplementary figure 3, we can see in A the evolution of the total error, function of the adaptation or synaptic strength variations around the selected values. We can see that the activity patterns properties are mostly sensitive to variations in PYR to PYR and PYR to VIP connection strengths, and PYR activity adaptation strength. All three parameters determine the changes in H-state durations, as seen in figure B, but the adaptation parameter is the only determinant in the changes of the mode of the distribution (see figure C).

In order to reproduce the variability in activity patterns observed in the WT mice, we simulated 100 networks, which parameters were determined from a Gaussian distribution, centered on the values obtained through the previously described selection procedure, with a standard deviation of 1 for synaptic weights and 0.1 for external inputs (∼5% of the initial values). We selected the parameter sets that reproduced most closely each mouse activity states duration properties. See supplementary figures 4 and 5 for H-state and L-state duration distributions in each recorded mouse, compared to simulations.

## Modeling nAChRs

We modelled the external inputs to the different neuronal populations as either cholinergic and regulated nicotinic receptors (I_β2-s_, I_α7-s_, I_α7-p_ and I_α5-v_), or non-cholinergic/non-regulated by nicotinic receptors (I_0-s_, I_0-p_ and I_0-v_). We used a minimal model of subtype-specific activation and sensitization of nAChRs [20], from which we determined the amplitude of each cholinergic current:

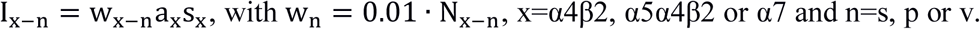

a_x_ is the activation variable, while s_x_ is the sensitization variable. They both take values between 0 and 1. The receptor is fully activated for a_x_=1 and fully sensitized for s_x_=1, while it is closed for a_x_=1 and fully desensitized for s_x_=0. We did not take into account desensitization of nAChRs by physiological levels of ACh because of its rapid breakdown through acetylcholinesterase [21, 22], such that s_x_=1. Using formulas from [20], with ACh=1.77 μM, relevant for in vivo simulations, we found a_α4β2_= a_α5α4β2_= 0.0487, and a_7_= 0.0014. N_n_ is the number of nAChRs of each type that best reproduced the changes of activity patterns recorded in experimental data (see tables 5 and Fig. 4A1-A2-A3). See table 6 for the corresponding cholinergic currents. For the nicotine application simulations, we used a physiologically relevant blood concentration for smokers Nic=1 μM and computed the change in sensitization and activation of the various nAChRs (see table 7). For α5α4β2 nAChRs, more resistant to desensitization, we shifted DC_50_ from 61 nM to 610 nM [24]. See table 8 for the corresponding cholinergic currents.

**Table 6:** Selected external currents (arbitrary unit).

| $I_{ext-e}$ | $I_{0-p}$ | $I_{\alpha 7-p}$ | $I_{0-s}$ | $I_{\alpha 7-s}$ | $I_{\beta 2-s}$ | $I_{0-v}$ | $I_{\alpha 5-v}$ |
| --- | --- | --- | --- | --- | --- | --- | --- |
| 3.9 | 4.4 | 0.1 | 2.1 | 0.1 | 1.4 | 2.55 | 0.35 |

**Table 7:** Activation and sensitization of nAChRs with Nic=1 μM.

| $a_{\alpha 7}$ | $s_{\alpha 7}$ | $a_{\alpha 4\beta 2}$ | $s_{\alpha 4\beta 2}$ | $a_{\alpha 5\alpha 4\beta 2}$ | $s_{\alpha 5\alpha 4\beta 2}$ |
| --- | --- | --- | --- | --- | --- |
| 0.0050 | 0.6282 | 0.1267 | 0.1981 | 0.1267 | 0.4385 |

**Table 8:** Cholinergic currents (arbitrary unit) with Nic=1 μM.

| $I_{\alpha 7-p}$ | $I_{\alpha 7-s}$ | $I_{\beta 2-s}$ | $I_{\alpha 5-v}$ |
| --- | --- | --- | --- |
| 0.23 | 0.23 | 0.732 | 0.405 |

In order to model the different KO mice, we set the corresponding cholinergic currents to 0. For the modelled α5 SNP mice, expressing a human polymorphism of the α5 nAChRs associated with both schizophrenia and heavy smoking, we assume that 30% of the α5 receptors on VIP are inactive.

## Modeling firing activity of single neurons

To see if we could reproduce neural activity distributions recorded from the various mutant mice [4], we simulated the firing rate of single neurons using the population model. You can observe in supplementary figure 6C the distribution of neural firing rates for two populations of simultaneously recorded cells in two different mice, which spiking activities are shown in A and B. Population 1 (in blue), shows lower levels of neural activities compared to population 2 (in red), but also lower activity variability between cells. Most of single cells mean activities are centered on the mean, located at the peak of the distribution. In population 2, we see that some neurons fire at much higher frequencies compared to most of the cells, creating a heavy tail in the distribution. Both distributions were well fitted by a log-normal function (see the blue and red lines), consistent with previous experimental observations [50]. In figure D, we can see the relation between the mean and the standard deviation of neural firing rates in WT mice populations of simultaneously recorded cells. It was well fitted by a linear function y=x. Interestingly, it has been suggested from previous computational modeling work that in cortical networks, skewed shape distribution of firing rates comes from the nonlinearity of the in vivo f-I curve [51, 52]. Hence, we simulated the firing rate of 50 single cells per population, using a log-normal distribution with mean equal to the simulated population firing rate, and such that its mean equals its standard deviation. We simulated the activity of 11 populations per mouse, for 200 sec. In order to model the activity of populations without L-states/H-states transitions, we used Gaussian noise around each parameter, with a standard deviation of 0.5 for synaptic weights and 0.05 for external inputs. This level of noise permitted to induce state transitions in 90% of simulated WT mice populations, close to the experimental value. All boxplots in Fig. 5 and 6 are exhibiting the distribution of mean neural activities between the 11 simulated mice, for a given mouse type, as in [4].

## Legends

**Supplementary figure 1: (A)** Distributions of ω_e-e_ (PYR to PYR), ω_p-e_ (PV to PYR) andω_s-e_ (SOM to PYR) synaptic weights parameter values, leading to networks with bistable dynamics (arbitrary units). Curves represent the probability distributions. The middle line shows the mean, the edge lines show the extremums. **(B)** Same as A, for ω_e-p_ (PYR to PV), ω_p-p_ (PV to PV) and ω_v-p_ (VIP to PV) synaptic weights. **(C)** Same as A, for ω_e-p_ (PYR to SOM) and ω_v-s_ (VIP to SOM) synaptic weights. **(D)** Same as A, for ω_e-v_ (PYR to VIP) and ω_s-v_ (SOM to VIP) synaptic weights.

**Supplementary figure 2: (A)** Distributions of ω_e-e_ (PYR to PYR), ω_p-e_ (PV to PYR) and ω_s-e_ (SOM to PYR) synaptic weights parameter values, for networks with bistable dynamics close to WT mice activity patterns (arbitrary units). **(B)** Same as A, for ω_e-p_ (PYR to PV), ω_p-p_ (PV to PV) and ω_v-p_ (VIP to PV) synaptic weights. **(C)** Same as A, for ω_e-p_ (PYR to SOM) and ω_v-s_ (VIP to SOM) synaptic weights. **(D)** Same as A, for ω_e-v_ (PYR to VIP) and ω_s-v_ (SOM to VIP) synaptic weights. **(E)** Distribution of the changes of high activity state levels for PYR, PV, SOM and VIP neurons, after KO of α5 receptors. The dashed line shows the reference at 0, for no change. **(D)** Distribution of the changes of low activity state for PYR, PV, SOM and VIP neurons, after KO of α5 receptors. The dashed line shows the reference at 0, for no change.

**Supplementary figure 3: (A)** Evolution of the total fitting error, as a function of variations in the synaptic and adaptation strengths (arbitrary units). **(B)** Evolution of the error for the high activity state mean duration, as a function of variations in the synaptic and adaptation strengths (arbitrary units). **(C)** Evolution of the error for the high activity state mode of durations, as a function of variations in the synaptic and adaptation strengths (arbitrary units).

**Supplementary figure 4:** Distribution (probability density function) of high activity state durations (in seconds), for simulations (red) and experiments (blue), recorded from mouse 1 to 11.

**Supplementary figure 5:** Distribution (probability density function) of low activity state durations (in seconds), for simulations (red) and experiments (blue), recorded from mouse 1 to 11.

**Supplementary figure 6: (A)** Spiking activity of a population of simultaneously recorded cells, called population 1. The y-axis shows the neuron ID. Recordings lasted ∼216 seconds. **(B)** Same as A, for another population, called population 2. **(C)** Distributions of neural firing rate activities for population 1 (in blue) and population 2 (in red), in spikes/min. The blue and red lines are fitted lognormal functions on each distribution. **(D)** Standard deviation (std) of neural firing rate activities as a function of the mean neural firing rate activities in WT mice populations of simultaneously recorded cells, in spikes/min (blue crosses). The black line shows the theoretical relation: mean of neural firing rates = std of neural firing rates.

**Supplementary figure 7: (A)** Example of a time varying population mean activity of pyramidal neurons, computed from the network model. The dashed red line delineates the threshold between high and low activity states (H-states and L-states, respectively). Red periods correspond to H-states and blue periods to L-states. **(B)** Same as A, for parvalbumin interneurons from the same network as in A. **(C)** Same as A, for somatostatin interneurons from the same network as in A. **(D)** Same as A, for VIP interneurons from the same network as in A.

**Supplementary figure 8: (A1)** Spike trains of a population of simultaneously recorded PV cells in a WT mouse, obtained through deconvolution of spontaneous Ca^2+^ transients [4]. **(A2)** Time varying population mean activity of the PV neurons shown in A1. The dashed red line delineates the threshold between high and low activity states (H-states and L-states, respectively). Red periods correspond to H-states and blue periods to L-states. See [6] for more info on the methods. **(B1)** Same as A1 for simultaneously recorded SOM cells in a WT mouse. **(B2)** Same as A2, for SOM cells. **(C1)** Same as A1 for simultaneously recorded VIP cells in a WT mouse. **(C2)** Same as A2, for VIP cells.

**Supplementary figure 9: (A1)** H-state (red line) and L-state (blue line) PV activities as a function of the external input current to PV interneurons population. The dashed green line shows the selected parameter value to reproduce WT mice neural dynamics. KO of nAChRs is associated with a decrease of external currents (black arrow). **(A2)** Same as A1, for SOM activities as a function of the external input current to PV interneurons population. **(A3)** Same as A1, for VIP activities as a function of the external input current to PV interneurons population. **(B1)** Same as A1, for PV activities as a function of the external input current to SOM interneurons population. **(B2)** Same as A1, for SOM activities as a function of the external input current to SOM interneurons population. **(B3)** Same as A1, for VIP activities as a function of the external input current to SOM interneurons population. **(C1)** Same as A1, for PV activities as a function of the external input current to VIP interneurons population. **(C2)** Same as A1, for SOM activities as a function of the external input current to VIP interneurons population. **(C3)** Same as A1, for VIP activities as a function of the external input current to VIP interneurons population.

**Supplementary figure 10: (A1)** Boxplots of H-state durations, for WT control mice and WT mice treated with 7 days of chronic nicotine application, experimental data described in [4]. WT mice treated with 7 days of chronic nicotine application exhibit higher H-state durations than WT control mice (Kruskal Wallis, P<0.001). **(A2)** Same as A1, for simulations. **(B)** Change of median SOM activity for WT mice under 7 days nicotine treatment, from simulations. SOM activity is significantly decreased (Kruskal Wallis, P<0.001).

